# Somatic reversion impacts MDS/AML evolution in the short telomere syndromes

**DOI:** 10.1101/2021.05.26.445858

**Authors:** Kristen E. Schratz, Valeriya Gaysinskaya, Zoe L. Cosner, Emily A. DeBoy, Zhimin Xiang, Laura Kasch-Semenza, Liliana Florea, Pali D. Shah, Mary Armanios

**Affiliations:** Departments of Oncology, Johns Hopkins University School of Medicine, Baltimore, MD 21287; Departments of Genetic Medicine, Johns Hopkins University School of Medicine, Baltimore, MD 21287; Departments of Medicine, Johns Hopkins University School of Medicine, Baltimore, MD 21287; Departments of Molecular Biology and Genetics, Johns Hopkins University School of Medicine, Baltimore, MD 21287; Telomere Center at Johns Hopkins, Johns Hopkins University School of Medicine, Baltimore, MD 21287; Sidney Kimmel Comprehensive Cancer Center, Johns Hopkins University School of Medicine, Baltimore, MD 21287

## Abstract

Inherited mutations in telomerase and other telomere maintenance genes manifest in the premature aging short telomere syndromes. Myelodysplastic syndromes and acute myeloid leukemia (MDS/AML) account for 75% of associated malignancies, but how these cancers overcome the germline telomere defect is unknown. We used targeted ultra-deep sequencing to detect candidate somatic reversion mutations hypothesizing they may promote MDS/AML evolution. While no controls carried somatic mutations in telomere maintenance genes (0 of 28), 29% of adults with germline telomere maintenance defects carried at least one (16 of 56, P<0.001). In addition to *TERT* promoter mutations which were present in 19%, we identified *POT1* and *TERF2IP* mutations in 13%. *POT1* mutations impaired telomere binding and in some cases were identical to those found in familial melanoma associated with longer telomere length. Exclusively in patients with germline defects in telomerase RNA (TR), we identified somatic mutations in nuclear RNA exosome genes, *RBM7, SKIV2L2*, and *DIS3*, where loss-of-function upregulates mature TR levels. Paradoxically, somatic reversion events were more prevalent in patients who were MDS/AML-free (P=0.01, RR 5.0, 95% CI 1.4-18.9), and no MDS/AML patient had more than one mutant clone (P=0.048). Our data identify diverse somatic adaptive mechanisms in the short telomere syndromes, and raise the possibility that their presence alleviates the telomere crisis that promotes transformation to MDS/AML.

---

Germline mutations in telomerase and other telomere-related genes are thought to be one of if not the most common known genetic cause of adult-onset myelodysplastic syndromes and possibly acute myeloid leukemia (MDS/AML)(1-4). In line with this observation, in patients with inherited mutations in telomerase and other telomere maintenance genes, MDS/AML are the most common cancers(5, 6). They account for 75% of short telomere syndrome malignancies but the lifetime risk is 10%(5, 6). The vast majority of short telomere syndrome MDS/AML cases are age-related; they manifest after the age of 50 often concurrently with idiopathic pulmonary fibrosis or other telomere-mediated pulmonary disease(5-8). In primary cells, critically short telomeres provoke a DNA damage response that triggers a p53-dependent checkpoint that signals apoptosis or cellular senescence(9, 10). This checkpoint mediates the degenerative phenotypes of the short telomere syndromes which manifest most prominently as bone marrow failure and pulmonary fibrosis(11, 12). How these myeloid cancers overcome the background short telomere checkpoint to sustain their replicative potential remains unknown. Moreover, the factors that predict which patients go on to develop MDS/AML are not understood. The latter question is particularly timely since pulmonary fibrosis patients with short telomere syndromes are now identifiable prior to lung transplantation, but have a significantly increased risk of acquiring post-transplant MDS and AML(6).

Here, we used an ultra-deep sequencing approach to test the hypothesis that acquired clonal somatic mutations that offset the inherited short telomere defect arise in telomere-mediated MDS/AML. As we show, in contrast to our expectations, we found diverse somatic reversion mechanisms that show great specificity relative to the germline defect. Surprisingly, however, these somatic mutations were overall rare in MDS/AML patients and had a higher prevalence and allele burdens in patients who were MDS/AML-free. These data have implications for fundamental understanding of how somatic adaptation in the hematopoietic system may provide protection against MDS/AML evolution in at least some inherited bone marrow failure syndromes.

## Results

To understand the mechanisms of telomere maintenance in short telomere MDS/AML patients, we compared the prevalence of candidate somatic mutations among 84 individuals who were over the age 40 and who were divided across three groups: healthy controls (mean age 64, n=28,), short telomere syndrome patients who did not have MDS/AML (mean age 60, n=40) and short telomere syndrome patients who fulfilled World Health Organization criteria for MDS/AML (mean age 56, n=16). Figure 1A summarizes their clinical characteristics. Healthy controls had normal telomere length, while the other two groups had similarly abnormal short length defects (Figure 1B, C). We used a customized ultra-deep error-corrected sequencing platform(13, 14), and designed our experiment to detect low frequency somatic mutations down to 0.5% variant allele frequency (VAF). Using this pipeline, we targeted variants in 16 candidate genes in which mutations could functionally promote telomere maintenance. Their functions were grouped in four categories: 1) promoters of the core telomerase components, telomerase reverse transcriptase and telomerase RNA, *TERT* and *TR*, 2) the six shelterin genes of which four are mutated in cancer-prone families hypothesized to have long telomere syndromes(15, 16), 3) six nuclear RNA exosome genes involved in *TR* processing and where loss-of-function has been shown to increase mature TR levels, and 4) *ATRX* and *DAXX*, known to be mutated in cancers that rely on alternative lengthening of telomeres (ALT) mechanisms (Supplementary Table 1).

**Figure 1.**
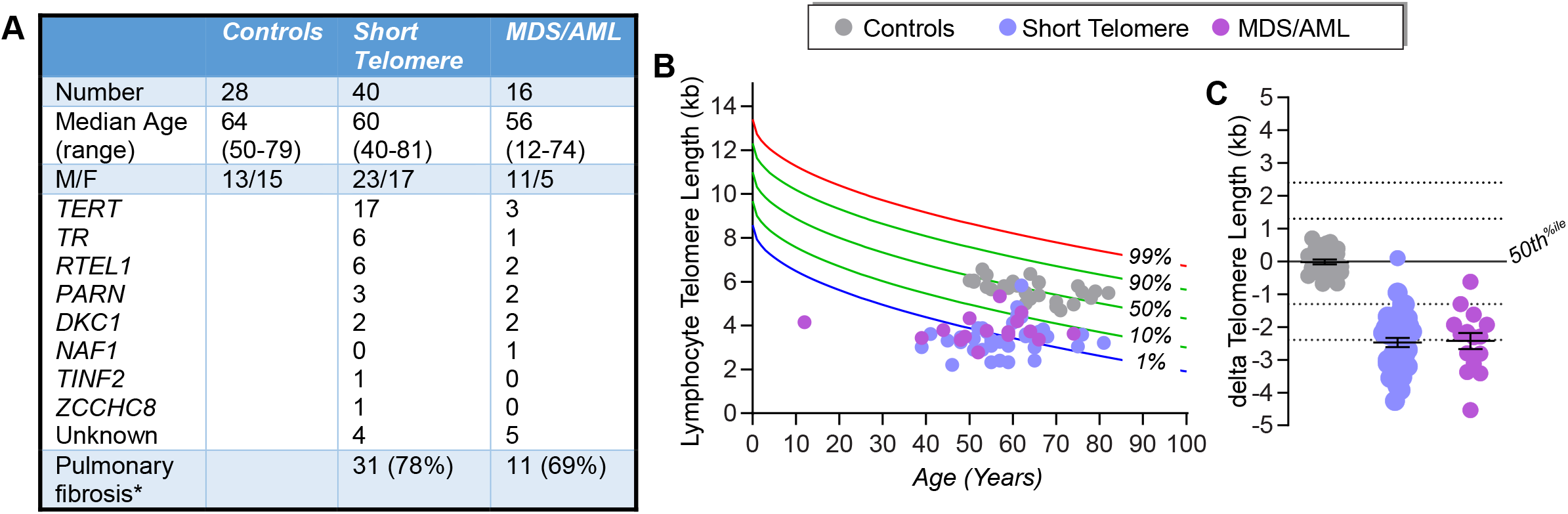
Clinical characteristics of controls, short telomere syndrome patients with and without myelodysplastic syndrome and acute myeloid leukemia (MDS/AML). **(A)** Demographics and mutant germline genes of controls and short telomere syndrome subjects. All the autosomal genes listed were heterozygous mutant. The unknown category included one patient with low telomerase RNA levels each. Pulmonary fibrosis refers to patients who had this concurrent diagnosis at the time of study recruitment and blood draw. **(B)** Telogram showing lymphocyte telomere length by flow cytometry and fluorescence in situ hybridization (flowFISH) relative to age-adjusted nomogram, and **(C)** is a plot of the difference of an individual lymphocyte telomere length relative to the median for age. Error bars represent standard error of the mean.

In addition, we examined *TP53* since its loss may allow bypass of the short telomere checkpoint(9). Among the three groups, we identified 36 clonal somatic mutations: 34 single nucleotide variants (SNVs) and 2 insertions/deletions. They fell in six of the 17 candidate genes (35%): *TP53*, the *TERT* promoter, *POT1, TERF2IP* and *RBM7, SKIV2L2* and *DIS3* (Figure 2A-D and Table 1). Only one was found in a healthy control; and that mutation was a low allele frequency *TP53* (VAF 1%) subclinical clonal hematopoiesis mutation. Another 10 *TP53* missense mutations fell predominantly in the DNA binding domain, and all are recurrently mutated in cancer (Figure 2A). The latter were divided equally among short telomere patients with and without MDS/AML (Figure 2A).

**Figure 2.**
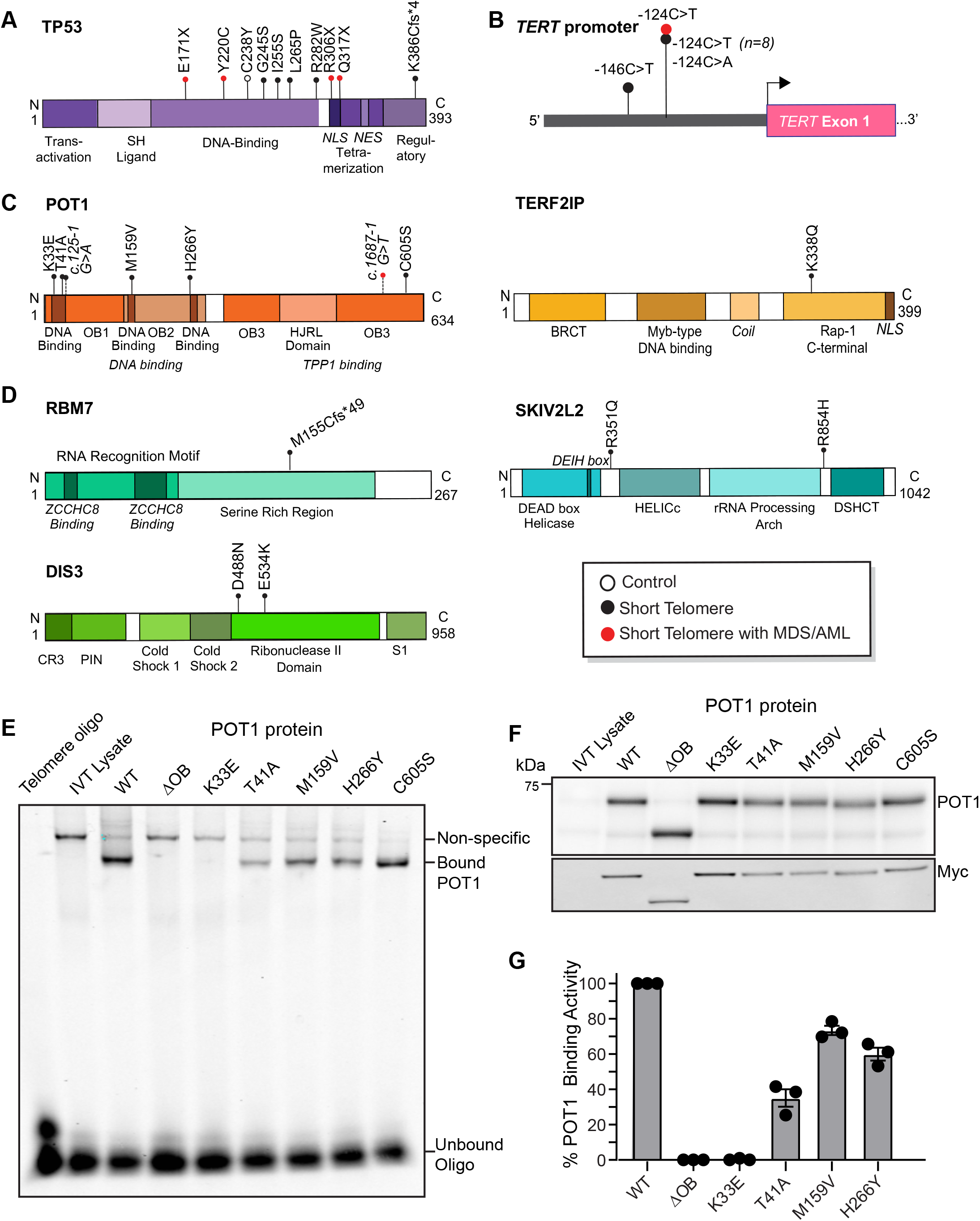
Somatic mutations relative to protein functional domains and POT1 functional studies. **(A), (B), (C)** and **(D)** Schema showing somatic mutations relative to mutant protein/gene with conserved domains drawn for TP53, the *TERT* promoter, shelterin components and nuclear RNA exosome components, respectively. The key denotes the group in which the mutation was identified with open circle referring to controls, black circles to short telomere patients without MDS/AML, and red circles to short telomere patients with MDS/AML. **(E)** Electrophoretic mobility shift assay (EMSA) for POT1 mutants examining the binding capacity of each of the missense mutations identified. POT1^OB^ refers to a control deleted for the first DNA binding domain (aa127-635). WT refers to *wild-type*. **(F)** Immunoblot of in vitro translated products shows missense mutant POT1 is stable with the Myc blot confirming specificity. **(G)** Mean intensity of binding relative to WT with error bars representing standard error of the mean. The data shown are from three EMSA reactions derived from two independent in vitro translations.

**Table 1.**
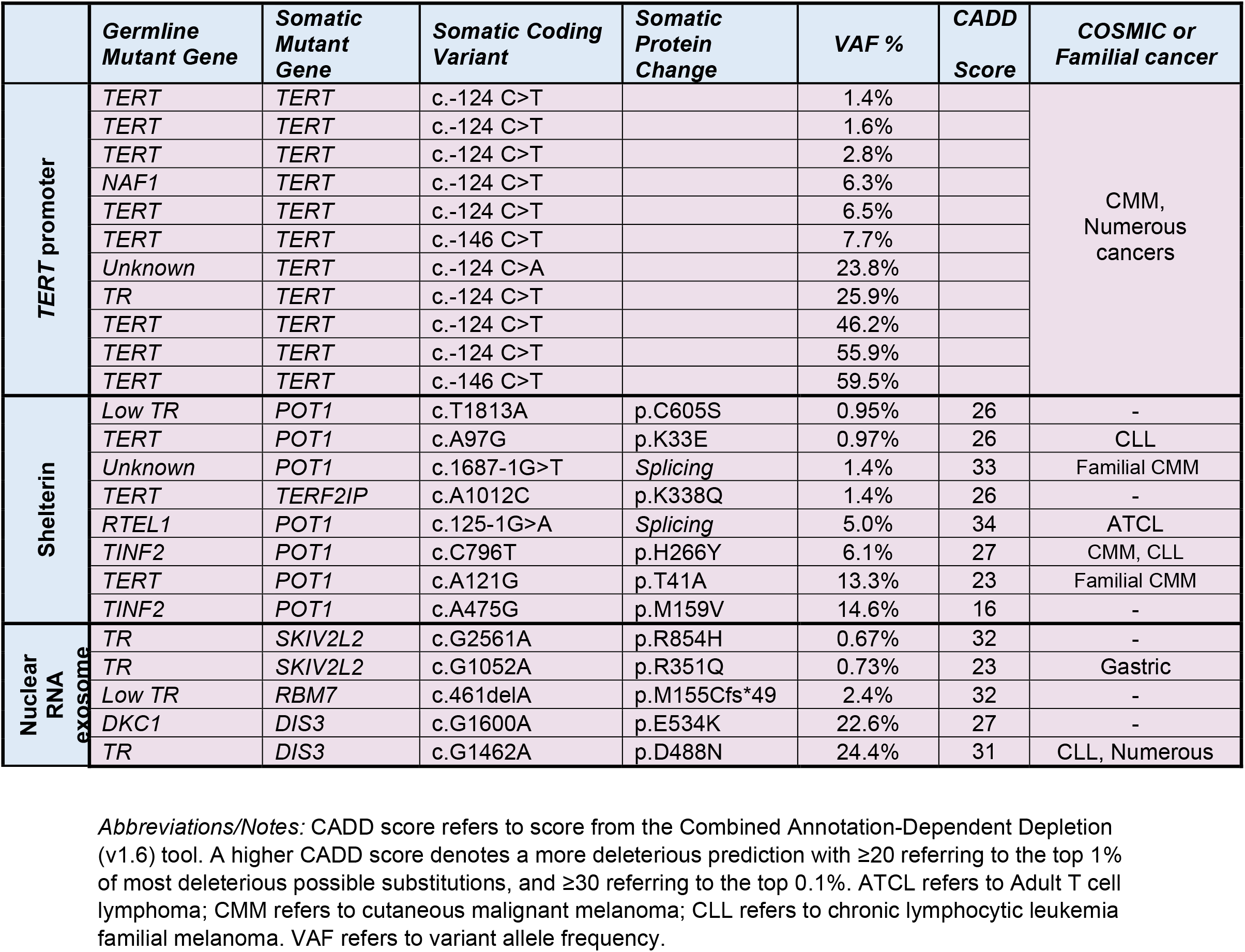
Germline and somatic mutations in telomere-related genes, their VAF and CADD score.

All the remaining somatic telomere gene mutations fell exclusively in short telomere syndrome patients with 30% carrying at least one (16 of 56 short telomere vs. 0 of 28 controls, P<0.001, Fisher’s exact test) (Table 1). The most common mutations were in the *TERT* promoter; they were seen at the -146 and -124 hotspots upstream of *TERT*’s transcription start site (Figure 2B). Nineteen percent (11 of 56) of the combined short telomere patients carried at least one of these mutation; this rate is higher than the 5-7% prevalence previously reported in patients with pulmonary fibrosis and dyskeratosis congenita which utilized less sensitive sequencing methods(17, 18). These *TERT* promoter mutations are identical to somatically mutated nucleotides found in numerous cancers and which create *de novo* ETS-binding sites that upregulate transcription of the intact *TERT* allele in *trans* with the germline mutation(17, 19-21).

The next most common class of mutations fell in *POT1*, a negative regulator of telomere elongation. Two mutations fell in canonical splice junction sequences (Figure 2C and Table 1). For *POT1*’s and the missense mutations in the other four protein-coding genes, all but one had Combined Annotation-Dependent Depletion (CADD) scores ≥20 signifying they represent the top 1^st^ percentile of damaging mutations, and five had a score of ≥30 indicating they were in the top 0.1 percentile of damaging mutations (Table 1). Several of the shelterin gene mutations, and mutations in nuclear RNA exosome genes, have been documented to be also somatically mutated in cutaneous malignant melanoma (CMM), chronic lymphocytic leukemia (CLL) among other cancers (Table 1). Somatic mutations at the POT1 p.H266 residue, for example, are the most common telomere maintenance mechanism mutated in CLL(22, 23), and two mutations, POT1 p.T41A and c.1687-1G>T have been identified in familial melanoma, a long telomere syndrome phenotype(15, 24) and DeBoy and Armanios, unpublished). A third group of mutations fell in RNA processing genes, and the impact of loss-of-function of the nuclear RNA exosome targeting (NEXT) complex genes, *RBM7, SKIV2L2*, and the nuclear RNA exosome essential nuclease, *DIS3*, on increasing TR levels has been previously documented(25, 26). Given the high frequency of *POT1* missense mutations, we tested their functional consequences in an electrophoretic mobility shift assay (EMSA). The four missense mutations in the OB folds fell within 3.5 Angstrom’s of telomeric DNA in the POT1-telomere crystal structure(23, 27) and showed impaired DNA binding to varying degrees (Figure 2E-G). Notably, the K33E mutation phenocopied a mutant that was deleted for the entire first OB fold of POT1 (Figure 2E-G). In contrast, POT1 p.C605S, which fell in the TPP1-interacting domain (Figure 1C), showed no DNA binding defects. These collective data supported that multiple *de novo* somatic mutations arise relatively commonly in telomerase and telomere gene mutation carriers. They are functionally impactful and overlap with somatic telomere maintenance mutations seen in cancer.

We next examined if the telomere-related somatic mutations clustered in the MDS/AML patients, but found a surprising and paradoxical result. Short telomere adults who were MDS/AML-free had more mutations than the MDS/AML patients (RR 5.0, 95% CI 1.4-18.9, P=0.01, Fisher’s exact test). Among adults without MDS/AML, 35% had ≥1 and 18% had ≥2 somatic mutations (14 and 7 of 40, respectively, Figure 3A). But for MDS/AML patients, only 13% (2 of 16) carried ≥1 and none carried ≥2 mutations (P=0.1 and P=0.08, respectively). The somatic mutant clones were also on average 3.8-fold larger in MDS/AML-free adults (mean 14.7% vs. 3.9%). Notably, none of the MDS/AML patients had a clonal somatic mutation in a telomere gene >10% VAF (P=0.048, Fisher’s exact test) (Figure 3B), suggesting this 10% threshold may be a useful predictive threshold if validated in other studies. We assessed the incidence of MDS/AML in the patients who carried somatic reversion mutations and all of them remained MDS/AML-free with a median follow-up of 2.0 years (range 3 months-11 years).

**Figure 3.**
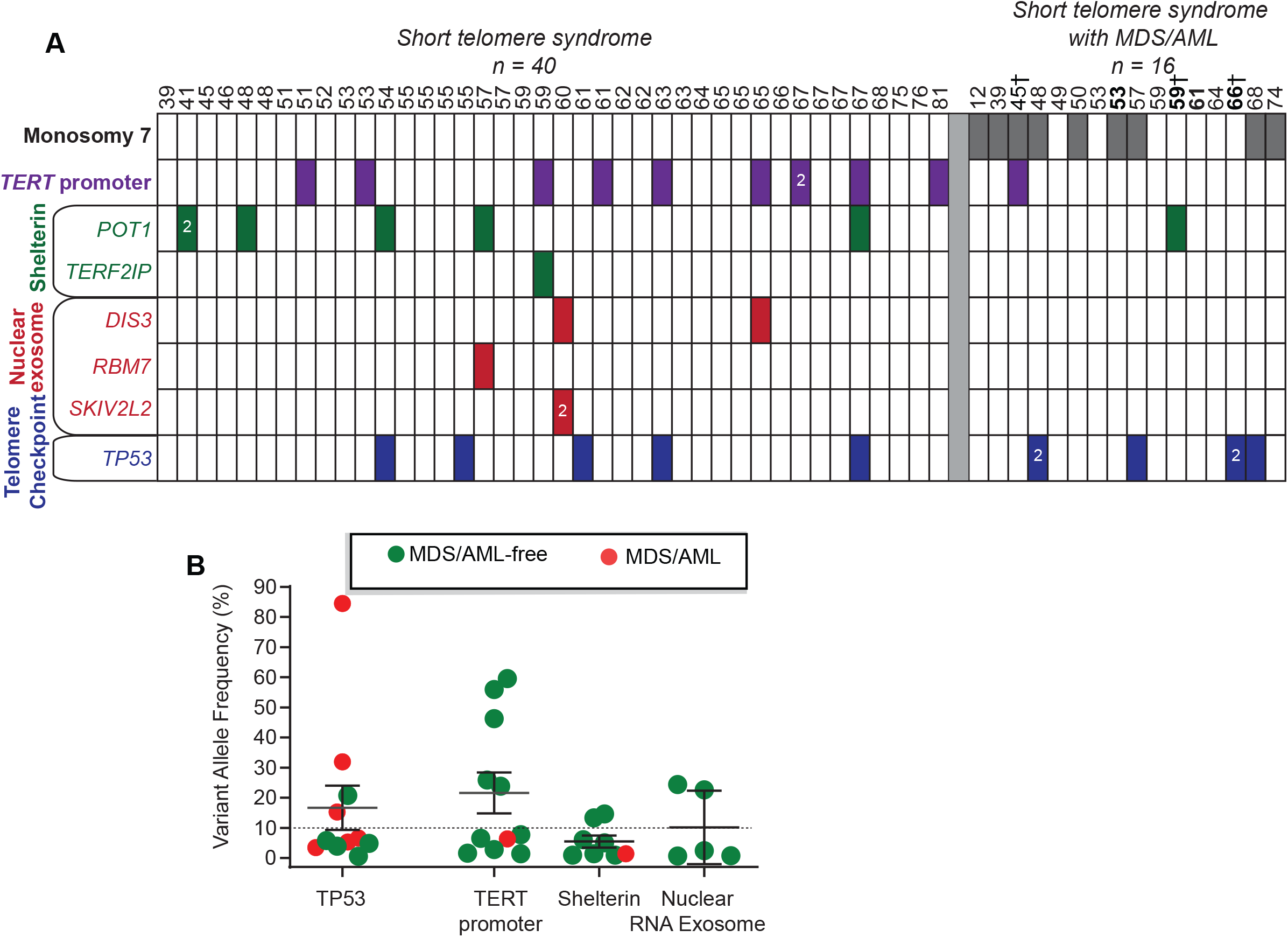
Prevalence of mutations relative to patient age and diagnosis of myelodysplastic syndrome or acute myeloid leukemia diagnosis (MDS/AML). **(A)** Chart shows individual patients with age noted above and the mutant gene identified in each column. The data are divided by presence or absence of MDS/AML. Monosomy 7 refers to patients who had any monosomy 7 abnormality including -7, del(7q) or der(1;7)(q10;p10)). Bolded ages refer to individuals for whom DNA was bone marrow-derived. †Denotes patients who were analyzed for the four common mutant genes on a clinical panel in contrast to others sequenced on the the customized panel. **(B)** Clone size plotted as variant allele (VAF) frequency by mutant gene or gene group. Shelterin refers to mutations in *POT1* and *TERF2IP/RAP1*, and Nuclear RNA Exosome refers to mutations in *RBM7, SKIV2L2* and *DIS3* as also labeled in **(A)**.

If these somatic mutations correlate negatively with MDS/AML onset, we would predict they would be limited to or enriched in the myeloid lineage. To test this directly, we used droplet digital PCR probes to quantify the mutant fraction in whole blood and compared it to myeloid and T cell fractions. We studied four mutations (*POT1* and *DIS3*) from three patients and found they were all enriched in the myeloid lineage while being low or undetectable in the T cell fraction (Figure 4). To definitively test if the somatic mutations we identified were indeed reversion events, we correlated the germline and somatic mutant pathways and found remarkable associations. While *TERT* promoter mutations predominated in germline *TERT* mutation carriers (8 of 11), *POT1* and *TERF2IP* mutations were associated with a more heterogeneous spectrum of germline mutations including *TERT, TINF2, RTEL1* and *PARN* (Table 1). This observation is consistent with POT1 loss-of-function mutations playing a role in improving telomerase access or processivity(15). By contrast, mutations in the nuclear RNA exosome genes were exclusively detected in patients with germline defects in TR abundance or function. All three patients who carried these somatic mutations carried germline mutations in *TR* itself, the ribonucleoprotein *DKC1*, or had abnormally low *TR* levels (Table 1). These data support a model where myeloid-specific adaptation that arises in the highly replicative environment of hematopoiesis, and under the selective pressures of a germline short telomere defect, is associated with an MDS/AML-free state.

**Figure 4.**
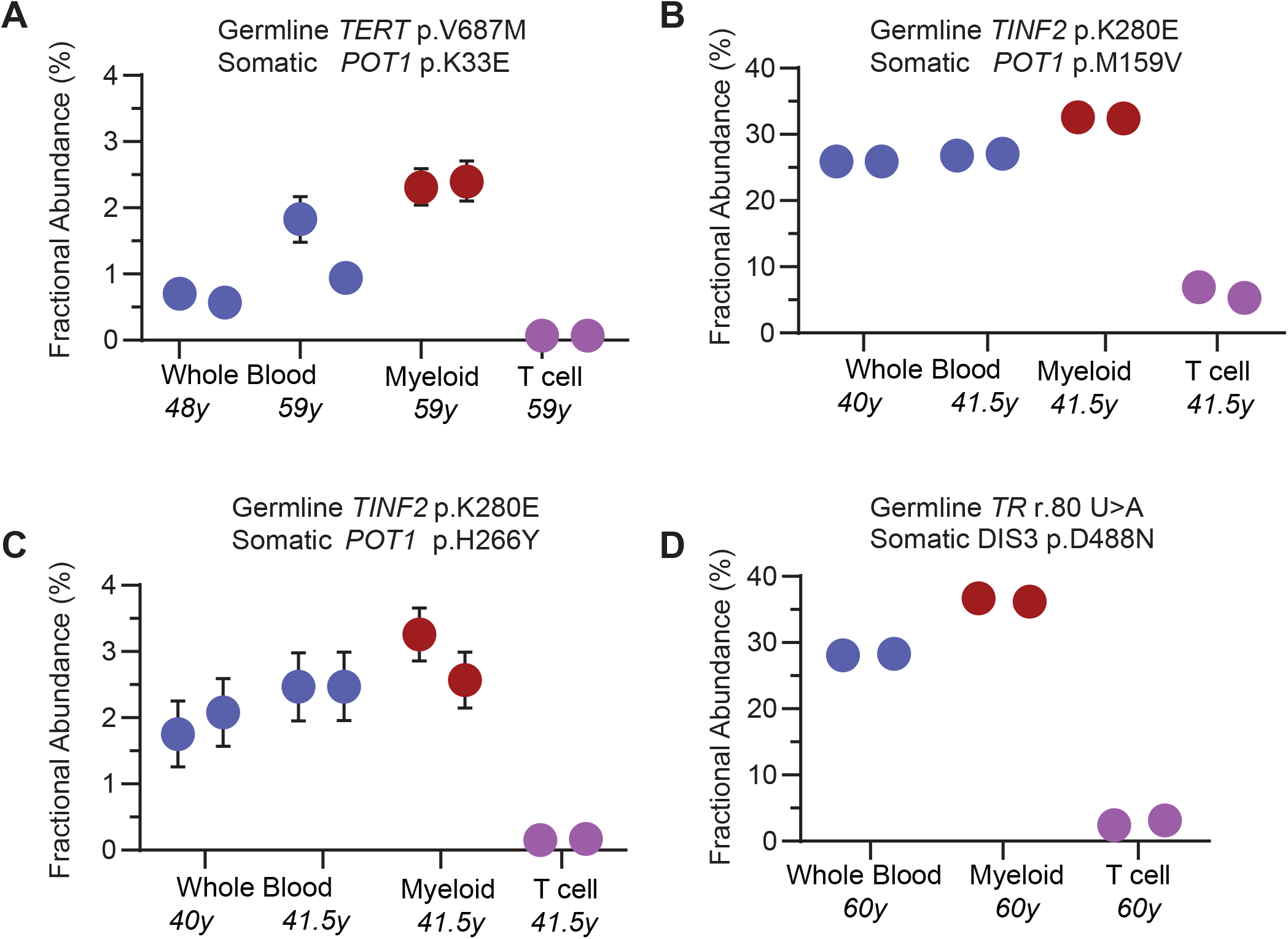
Fractional abundance of mutant clone size in leukocyte fractions quantified by droplet digital PCR (ddPCR). **(A-D)** ddPCR abundance of mutations are shown at study enrollment, and for three of the mutations at follow-up. The germline and somatic mutations are shown above, and the age at the time of draw is annotated below. DNA for ddPCR analysis was derived from whole blood, the myeloid fraction (CD33^+^ CD66^+^) and CD3^+^ T cell fraction. Each dot refers to mean fractional abundance from the Poisson distribution with error bars representing the minimum and maximum of the dataset with each reaction performed twice denoted. Where not seen, min/max lines fall within the boundaries of the graphed dot.

## Discussion

In the high turnover environment of hematopoiesis, the selective pressures imposed by inherited mutations allow advantageous clonal mutations to persist(28). Nearly all the known somatic reversion mechanisms are direct in that they repair the germline mutant gene itself such as by correcting a frameshift or through homologous recombination with the *wild-type* allele(28-30). Here, we identified several indirect somatic reversion mechanisms in the short telomere syndromes, and show they overlap with known telomere maintenance mechanisms in cancer.

Their high specificity, such as is the case for the nuclear exosome components in *TR* mutation carriers, underscores a remarkable versatility of the stressed hematopoietic system to acquire *de novo* adaptations. We did not observe gross measurable telomere lengthening effects (Supplementary Figure 1A), but recurrent mutations, often in the same genes or pathways within a given individual patient, highlight the competitive advantage they provide under the selective pressures of the short telomere environment. The finding of *POT1* recurrent mutations across multiple germline defects in *TERT* and other mutation carriers support a model where *POT1* loss-of-function facilitates telomere elongation by improving telomerase access and/or processivity. Of note, several of the mutations we identified have been seen in cancer prone families with melanoma. The shared spectrum of mutations we describe here with somatic mutations seen in cancer as well as in germline mutant cancer prone families support a model wherein *POT1* mutations themselves do not promote genome instability but instead favor a state of telomere maintenance(15).

To our surprise, the adaptive mechanisms we identified, including mutations in the *TERT* promoter, had a higher prevalence in patients who did not have MDS/AML. These data shed light into how the short telomere bone marrow failure state may evolve to myeloid malignancy. One possibility is that short telomere progenitors undergoing crisis undergo fate-determining events that lead to a functional reversion or a maladaptive mechanism such as loss of chromosome 7. In the majority of cases in our study, these events were mutually exclusive.

Comparing the 33% prevalence of reversion mutations we identified in adult short telomere patients (Figure 4), relative to the 10% overall incidence of MDS/AML, somatic reversion appears to be a more common and preferred adaptive mechanism. It is also possible that somatic reversion mutations may be similarly associated with an MDS/AML-free state in other MDS/AML inherited bone marrow failure states. Our data raise the possibility that deep sequencing for at least some of the common mutations we identified may be a useful biomarker for assessing MDS/AML risk in some settings. IPF is the most common indication of lung transplantation world-wide and it is estimated that 35% of familial pulmonary fibrosis patients carry a germline defect in a telomere maintenance gene(31, 32). In the setting of lung transplantation, short telomere patients are exposed to cytotoxic medications that add selective pressures on hematopoiesis, and the presence of sizable clonal reversion mutations may provide a protective advantage. The median follow-up for patients with telomere-related somatic reversion mutations was two years. Additional studies and longer follow-up will be needed to test the predictive utility of somatic reversion testing in clinical settings.

## Materials and Methods

### Subjects and study approval

Subjects were recruited 2003 to 2020 as part of the Johns Hopkins Telomere Syndrome Study(6, 33). They were enrolled if they carried a pathogenic mutation in a telomere-related gene *and* had a personal/family history of a short telomere phenotype, had classic familial short telomere syndrome as defined(7), *or* had sporadic disease with low telomerase RNA levels(34). The MDS/AML diagnosis was assigned using WHO classification(35). Healthy controls were recruited as previously and were selected for having normal telomere length(36). The research was approved by Johns Hopkins Medicine’s Institutional Review Board, and subjects gave written informed consent.

### Germline sequencing, functional studies and telomere length measurement

Germline sequencing was performed on peripheral blood DNA using whole genome, exome, or targeted sequencing(25, 30, 36). Germline mutations not reported in Alder *et al*.(36) or Schratz *et al*.(6) are listed in Supplementary Table 1. Telomerase RNA quantification and *DKC1* functional analyses have been detailed(25, 37). Telomere length was measured by flow cytometry and fluorescence in situ hybridization (flowFISH) as described(36, 38).

### Capture Design, Library preparation and Sequencing

We designed 150 bp read length probes (SureDesign) targeting 37 kb across 17 genes including two promoters (Supplementary Table 2). Libraries were prepared following the manufacturer’s instructions (Agilent Haloplex^HS^ for 1-500kb target region). Briefly, 400 ng of genomic DNA diluted to 14.4 ng/uL were digested in eight restriction enzyme reactions and hybridized to the custom biotinylated HaloPlex^HS^ probe library incorporating sample indexes and a degenerate 10-nucleotide unique molecular barcode (i.e. UMI, to identify each individual captured DNA molecule) to facilitate base calling accuracy and quantification. Targets were ligated to form circularized fragments, captured using streptavidin beads and polymerase chain reaction amplified (24 cycles). Target libraries were validated using Fragment Analyzer followed by quantification of fragment smear analysis of 190bp to 640bp range. All samples were subsequently pooled at 4 nM concentration. A final AMPure XP magnetic bead cleanup was performed on the pooled library to eliminate any adapter-dimer traces.

### Haloplex library sequencing

Sequencing was performed on an Illumina HiSeq 2500 using a 150 bp paired-end protocol, and data were demultiplexed following standard protocols. Reads were trimmed and aligned to the reference hg19 using BWA MEM in the Agilent Alissa Align and Call v.1.1.2.2 platform. Of 86 samples designed for Haloplex platform sequencing, 80 eventually fulfilled quality control criteria (3 failed at library preparation, 3 had low coverage). The mean depth of coverage was 12,522x ± 4,591 SD with a median of 10,952x ± 5,959 SD. The coverage depth for the *TERT* promoter was relatively lower and was calculated separately using Samtools (version 1.10, using htslib 1.10). Within the targeted interval (chr5:1295085-1295385, hg19), its coverage was limited to (ch5:1295085-1295115, hg19) and (chr5:1295224-1295385) and these intervals had a mean coverage of 2,457x ± 1,204 SD with a median 2,421x. Three additional MDS/AML patients were consecutively recruited after the Haloplex sequencing protocol started and their DNA was sequenced using a clinically validated dual-indexed targeted method that applied a background error rate analysis to allow robust detection of somatic mutations with allele frequency ≥1%(6, 39). This pipeline included the 3 most commonly mutated genes in the Haloplex analysis: *TP53, TERT promoter, POT1*, as well as *TINF2*. Supplementary Figure 1B lists the mutation types at the base pair level and shows 58% of SNVs were cytosine to thymidine transitions.

### Variant calling and analysis

Variant calling was performed using Agilent Alissa Align and Call v1.1.2.2 software and annotated using ANNOVAR v4162018. To utilize the Haloplex error-corrected method of somatic variation, we first collapsed reads originating from the same sample molecule into read families and duplicates were removed to create consensus reads. Variants were filtered to include only those present in ≥3 unique read families. To exclude sequencing artifacts, we used a quality score of >90 and filtered for a variant score threshold of >0.3 in Alissa. Variants that were present in >5% of controls were additionally excluded as alignment artifacts. Variants with allele frequencies of 40-60% or >95% were deemed germline and excluded. We finally filtered for variants with minor allele frequencies >0.0005 in gnomAD v.2.1.1 (any VAF), and these were excluded as they were thought to be possibly germline (performed November 1, 2020). The *TERT* promoter was manually analyzed and curated separately to include variants present in all reads of one or more consensus families at the canonical c.-124 and c.-146 positions irrespective of variant allele frequency. All of the variants reported were manually verified in Integrative Genomics Viewer (IGV).

### Variant interpretation

Protein schema were drawn based on the longest isoform curated in the Gene Database (National Center for Biotechnology Information, accessed 10/21/20). Linear protein structures were visualized in SnapGene (v5.1.6), and conserved domains constructed were based on Uniprot (release 2020_05) and the literature. Variants were analyzed for their presence in the COSMIC database (v.92) and Combined Annotation Dependent Depletion (CADD) score (v1.6)(40).

### POT1 gel shift binding assay

To test the functional significance of POT1 missense mutations, we cloned Myc-FLAG tagged POT1 into T7 expression vector (pcDNA 5/FRT, Addgene) and performed site-directed mutagenesis. A control with POT1^ΔOB^ was generated by deleting the first OB fold (amino acids 127-635). T7 expression vectors were used in an in vitro translation reaction using TNT Quick Coupled Transcription/Translation System (Promega), according to manufacturer’s instructions. Briefly, a 50 µl reaction containing 1 µg of expression vector, 20 µM methionine, and 40 µl TNT Quick Master mix containing rabbit reticulocyte lysate was incubated at 30°C for 90 minutes and used immediately for the gel shift assay.

The POT1 binding assay were preformed using Odyssey Infrared EMSA Kit (LI-COR). Briefly, a 20 µl reaction mixture was prepared containing 2 µl of the IVT product, 10 nM IRDye800-labled single stranded telomeric oligonucleotide, 2.5 µM nonspecific ssDNA and 1 µg poly(dI-dC), in binding buffer containing 100 mM Tris, 500 mM KCl, 12.5 mM DTT, 10 mM EDTA, and 0.25% Tween 20. We used this telomere oligonucleotide, 5’GGTTAGGGTTAGGGTTAGGG, and the non-specific ssDNA oligonucleotide 5’TTAATTAACCCGGGGATCCGGCTTGATCAACGAATGATCC as per Bauman *et al*.(41). Reactions were incubated for 30 minutes at room temperature and protein-DNA complexes were resolved by gel electrophoresis on a 10% polyacrylamide Tris-Borate EDTA at 100V for 2h at 4°C. Gels were visualized using Li-Cor Odyssey Imaging System. Quantification was done using ImageJ (2.1.0).

### Western blot

Using the NuPAGE SDS-PAGE gel system, 1 µl of in vitro translation reaction was run on 8% Bis-Tris gels with MOPS-SDS running buffer at 150 V and transferred to a PVDF membrane using the iBlot 2 Dry Blotting System on setting P0 (Thermo Fisher Scientific). Membranes were blocked for 1h before antibody staining. Membranes were stained first with anti-POT1 antibody (rabbit, NB500-176, 1:500, Novus) and subsequently with anti-rabbit horseradish peroxidase secondary antibody (HRP, Goat, 1:20,000, Cell Signaling Technology) before visualization via chemiluminescence (SuperSignal West Pico Plus Chemiluminescent substrate kit) using ImageQuant. Membranes were then stripped for 15 minutes, using New Blot IR Stripping buffer, and blocked again for 1h before staining with anti-Myc antibody (mouse, clone 4A6, 1:1000; Millipore). Membranes were subsequently stained with anti-mouse IRDye secondary antibody as appropriate (IR680, donkey, 1:10,000, LI-COR) before visualization on an Odyssey scanner (LI-COR). Quantification was done using ImageJ (2.1.0).

### Cell fractionation

Aliquots of fresh blood or frozen ficoll-separated peripheral blood cells were fractionated using EasySep Human Myeloid Positive Selection and EasySep Human T Cell Enrichment kits (StemCell Technologies. DNA was extracted from the fractionated subsets using MagMAX DNA Multi-Sample Ultra 2.0 Kit (ThermoFisher) and quantified using Quibit dsDNA HS Assay kit (ThermoFisher).

### Droplet Digital PCR

Droplet digital PCR (ddPCR) was performed according to manufacturer’s specifications on a QX200 system (Bio-Rad). Sequence-specific primer and probes were designed using the Bio-Rad digital assay design engine: *POT1* p.K33E (dHsaMDS726207399), *POT1* p.M159V (dHsaMDS784505022), *POT1* p.H266Y (dHsaMDS401027863), *DIS3* p.D488N (HsaMDS2512362). An annealing temperature of 58°C was used except for the *POT1* p.K33E (54°C) and *DIS3* p.D488N (54°C). The wildtype and mutant probes were conjugated with HEX and FAM reporters, respectively. PCR amplicons were sequence-verified by cloning the products (TOPO TA Cloning Kit, Invitrogen). For each target, the mean number of droplets analyzed was 11,890 ± 2,081 SD and all reactions had >9,000 events analyzed. Data were analyzed in QX Manager (version 1.2-STD, Bio-Rad).

## Statistics

Graphs were generated and statistical calculations were performed using GraphPad Prism 8.0.

## Author contributions

KES designed the study and analyzed the clinical and genetic data. KES and VG selected the candidate genes. VG prepared the libraries with LKS. ZLC performed the POT1 functional studies and analyzed ddPCR data. EAD did computational analyses. ZX performed the fractionation and ddPCR experiments. LF supported designing the sequencing analysis pipeline. PDS analyzed clinical data. MA oversaw the project and wrote the paper with KES. All the authors reviewed and approved the manuscript.

## Acknowledgements

We would like to acknowledge helpful conversations Dustin Gable, and technical support from the Johns Hopkins Genomics team, the Johns Hopkins Genetics Resources Core Facility and the Bloomberg School of Public Health Flow Cytometry Core. We are grateful for technical support from the Agilent team of Josh Zhiyong Wang, Gregory Miles and Weiwei Liu. This work was supported by RO1CA225027 and RO1HL119476, the Commonwealth Foundation, S&R Foundation Kuno Award, and the Williams Foundation (MA), T32HL007525, a grant from the Vera and Joseph Dresner Foundation and MacMillan Pathway to Independence Award (KES), F32HL142207 (VG), the Research Program for Medical Students (ZLC), T32GM007309 and the Turock Scholars Fund to the Telomere Center at Johns Hopkins (KES, EAD). The authors would also like to acknowledge a gift in the name of Mrs. P. Godrej.

## Conflict of Interest

The authors declare no conflicts of interest.

## Notes

### Competing Interest Statement

The authors have declared no competing interest.

