## Supplementary Materials for "Somatic reversion impacts MDS/AML evolution in the short telomere syndromes"

**Supplementary Table 1. Genes and regions (n=17) targeted for Haloplex sequencing**

| Gene Name | NM_ID/Coordinates |
|---|---|
| ***Telomerase Core Promoters*** | |
| ***TERT promoter*** | chr5:1295085-1295385 |
| ***TR promoter and TR gene*** | chr3:169482378-169483348 |
| ***Shelterin*** | |
| ***POT1*** | NM_015450 |
| ***TINF2*** | NM_001099274 |
| ***TERF2IP*** | NM_018975 |
| ***ACD/TPP1*** | NM_001082486 |
| ***TERF1*** | NM_017489 |
| ***TERF2*** | NM_005652 |
| ***RNA Exosome and RNA Processing*** | |
| ***MTREX/SKIV2L2*** | NM_015360 |
| ***RBM7*** | NM_001286045 |
| ***DIS3*** | NM_014953 |
| ***PAPBN1*** | NM_004643 |
| ***TENT4B/PAPD5*** | NM_001040284 |
| ***ZC3H18*** | NM_001294340 |
| ***ALT*** | |
| ***ATRX*** | NM_000489 |
| ***DAXX*** | NM_001141970 |
| ***Cell Cycle Checkpoint*** | |
| ***TP53*** | NM_000546 |

**Supplementary Table 2. Germline mutations in telomere or telomerase genes not previously reported***

| Mutant Gene | Coding Variant | Protein | Prior report |
|---|---|---|---|
| ***DKC1*** | c.942G>A | p.Lys314Lys | Gaysinskaya 2020[1] |
| | c.915+10G>A** | - | - |
| ***NAF1*** | c.950A>T | p.Asp317Val | - |
| ***PARN*** | c.543_544insTT | p.Asp182Leufs*5 | - |
| | NC_000016.9:g.(?_14725823)_(14643928_?)del | Exon 1-21 deletion*** | Feurstein 2020[2] |
| | c.(?_-135_(c.1480+1_1481-1)del | Exon 1-21 deletion*** | - |
| ***TERT*** | c.345C>G | p.Phe115Leu | - |

*Remaining mutations have been reported in Alder *et al.* 2018 and Schratz *et al.* 2020[3, 4].

**This mutation was identified in association with a more common *RTEL1* variant of unknown significance but because of rarity, was felt to be the more likely culprit.

***These two individuals shared a deletion of the *PARN* exons but it is unclear whether they shared the same breakpoints.

### Supplementary Figure 1

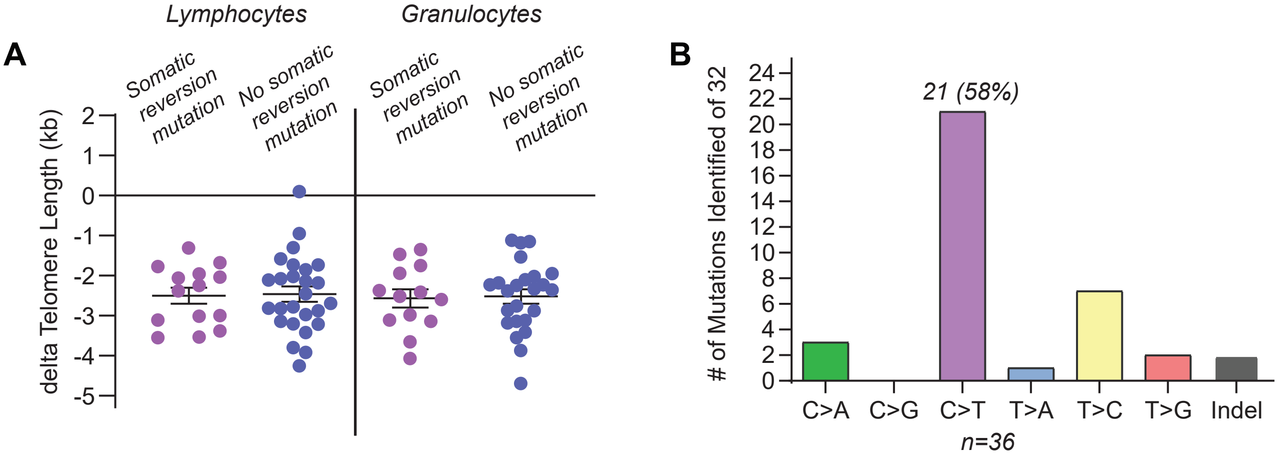


**Mutation types detected and their clone size and effect on telomere length. A.** Comparison of telomere length graphed as the difference from the age-adjusted 50th percentile in lymphocytes and granulocytes. Data are stratified by the presence or absence of a somatic reversion telomere-related mutation. The mean is shown and error bars refer to standard error of the mean. **A.** Mutation types identified in cohorts (controls, short telomere and short telomere with MDS/AML).
